# When and where to count? Implications of migratory connectivity and non-breeding distribution to population censuses in a migratory bird population

**DOI:** 10.1101/2022.02.26.482079

**Authors:** Antti Piironen, Anthony D. Fox, Hakon Kampe-Persson, Ulf Skyllberg, Ole Roland Therkildsen, Toni Laaksonen

## Abstract

Migratory connectivity is a metric of the co-occurrence of migratory animals originating from different breeding sites, and like their spatio-temporal distributions, can vary substantially during the annual cycle. Together, both these properties affect the optimal times and sites of population censusing.

We tracked taiga bean geese *(Anser fabalis fabalis)* during 2014–2021 to study their migratory connectivity and non-breeding movements, and determine optimal periods to assess the size of their main flyway population. We also compared available census data with tracking data, to examine how well two existing censuses covered the population.

Daily Mantel’s correlation between breeding and non-breeding sites lay between 0 and 0.5 during most of the non-breeding season, implying birds from different breeding areas were not strongly separated other times in the annual cycle. However, the connectivity was higher among birds from the westernmost breeding areas compared to the birds breeding elsewhere. Daily Minimum Convex Polygons showed tracked birds were highly aggregated at census times, confirming their utility. The number of tracked birds absent at count sites during the censuses however exceeded numbers double-counted at several sites, indicating that censuses might have underestimated the true population size.

Our results show that connectivity can vary in different times during the non-breeding period, and should be studied throughout the annual cycle. Our results also confirm previous studies, which have found that estimates using marked individuals usually produce higher population size estimates than total counts. This should be considered when using total counts to assess population sizes in the future.

## 1. Background

Reliable, accurate and regular population size estimates are essential for evaluating the conservation status of populations (Maes et al., 2015), setting targets for management and assessing the impact of population management actions (Madsen et al., 2017). To assess sizes of migratory populations or sub-populations, knowledge about the degree of migratory connectivity (Webster et al., 2002) throughout the annual cycle is essential. Migratory connectivity determines the co-occurrence of birds originating from different breeding sites throughout the annual cycle. This property is high when individuals from same breeding populations remain close throughout their annual cycle and separate from those of other breeding populations, whereas it is low when individuals remain close at one stage of the annual cycle but not at another, so providing a useful measure of how separate elements of a population may remain throughout the annual cycle (Webster et al., 2002; Cohen et al., 2017).

The strength of migratory connectivity between breeding and non-breeding sites can vary between different phases of the non-breeding seasons (Knight et al., 2021). Measurements of connectivity help to reveal clustering of the population through the non-breeding season and its implications for population size assessment (i.e. when and where individuals should be counted to avoid missing any clusters). Similarly, spatio-temporal distributions of the migratory populations can vary substantially during the annual cycle, which has obvious implications for when and where population censuses should optimally be done. Together, measurements of connectivity and spatial dispersion of populations over the annual cycle help identify the most favourable periods for population censusing. Although modern tracking technology provides efficient tools to study these pre-requisites for population censuses, we are not aware of any such studies (but see Finger et al., 2016 for study comparing timing of spring migration and breeding bird monitoring).

A variety of methods have been developed to monitor waterbird populations (Delany & Scott, 2005), but the assessment of goose population sizes is usually based on so-called total counts (Fox & Leafloor, 2018). These counts are often undertaken in mid-winter, when geese are most highly aggregated and when turnover of individuals, more likely associated with migratory staging areas, is considered to be at its lowest. During these counts, birds are censused at as many known different sites as possible (usually during a short period of time) and the population size is estimated as a sum of birds counted from different sites. These counts are based on the assumption that only a negligible amount of birds are missed in the counts (i.e. all birds are found) or are double-counted (i.e. birds have not moved between count sites during the count). The performance of these schemes are seldom evaluated, although some comparisons with capture-mark-resight estimates (Ganter & Madsen, 2001; Clausen et al., 2019) and model predictions (Johnson et al., 2020) have been made.

In contrast to several other goose populations throughout the globe, the Western Palearctic population of taiga bean goose (*Anser fabalis fabali*s, hereafter taiga bean goose) has declined throughout its range in recent decades (Fox & Leafloor, 2018). The whole population of the subspecies has recently been divided into four flyway populations (or management units, Marjakangas et al. 2015; Heinicke et al., 2018). The main flyway for the taiga bean goose is the Central Flyway (hereafter CF), which breeds in Finland, Sweden, Norway and North-Western Russia (Heinicke et al., 2018; see also Figure 1). The majority of the CF population is thought to winter in southern Sweden (e.g. Nilsson, 2011), but migration patterns and wintering sites of the birds breeding in North-Western Russia remain unknown. Additionally, taiga bean geese, thought to be from this flyway (Nilsson et al., 1999), winter in Denmark and northern Germany (Heinicke et al., 2018), but their origin and migration patterns are largely unknown (but see Nilsson, 2011; Mitchell et al., 2016; Boer, 2019 for some insights).

**FIGURE 1.**
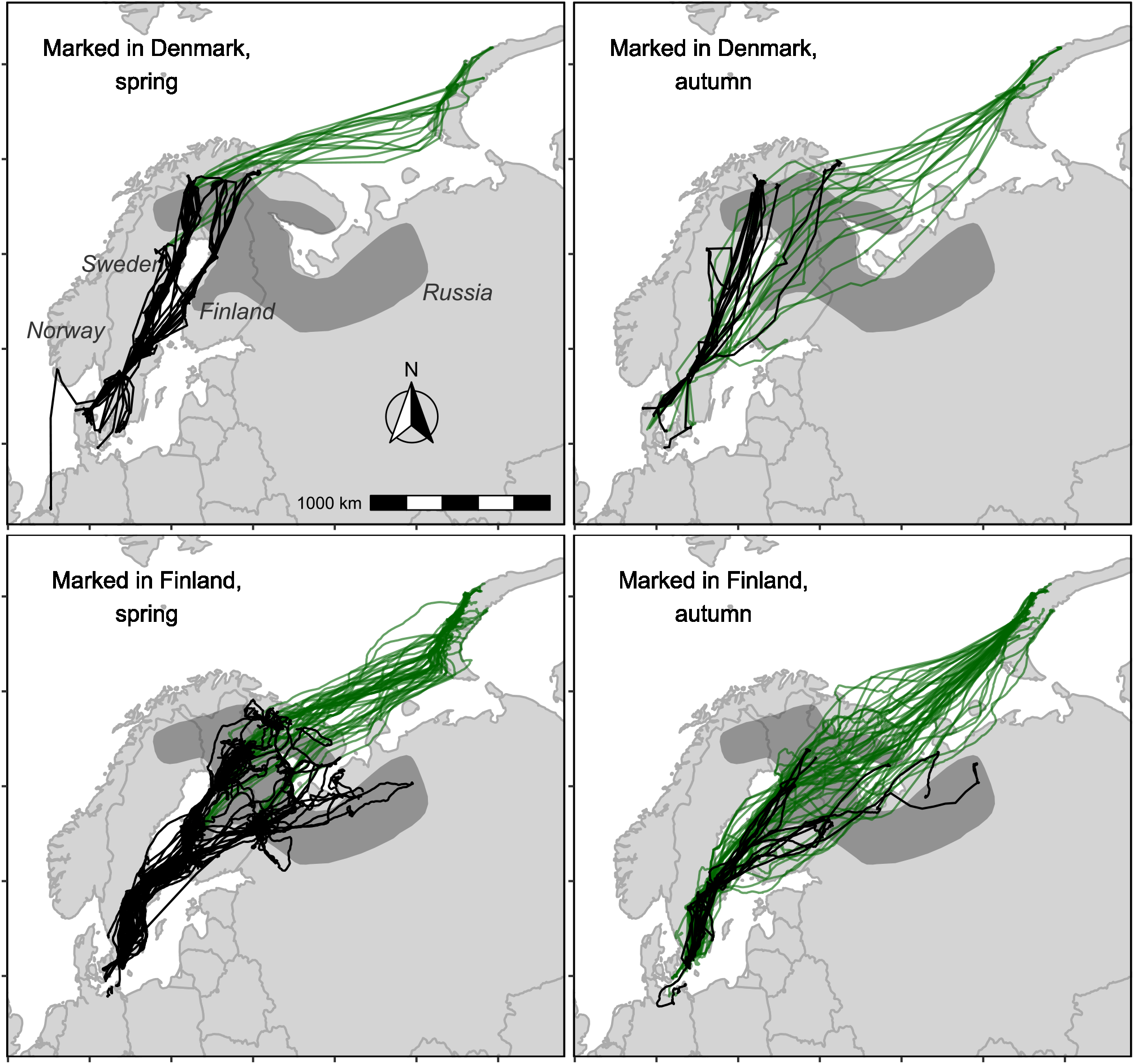
Migration routes of taiga bean geese marked for satellite tracking in Denmark in the winter (maps on the top row) and in Finland in the spring (maps on the bottom row). Figure shows all data from all tracked individuals (n = 68) from the years 2015–2021. Maps showing the spring and autumn migration routes include locations from the periods 1 January–30 May and 1 August– 31 December, respectively. To ensure figure clarity, migration routes to moulting sites at Novaya Zemlya (1 June–31 July) and back to wintering sites (1 August–31 December) are illustrated by green lines, while black traces show spring and autumn routes taken to and from the breeding sites (i.e. not moult migrants).

Population size assessment is highly relevant for the international adaptive harvest management of the CF population, since a target size for the population is set to 60 000–80 000 individuals (Marjakangas et al., 2015; Johnson et al., 2016). At the start of the flyway-scale management of the population, it was agreed to use mid-January counts to monitor the CF population (Marjakangas et al., 2015). In addition to the mid-winter counts, large-scale, coordinated counts of taiga bean geese were carried out in Swedish staging areas in October (autumn counts; see Nilsson & Kampe-Persson, 2020) and March (spring counts; see Skyllberg, 2015). It was suspected (but never verified) that at these times the vast majority of the flyway population was present, because these spring and autumn counts always far exceeded those counted in mid-winter (Johnson et al., 2021). Currently, estimates generated by the integrated population model are used to monitor the status of the population, using data from October, mid-winter and March as inputs in the model (Johnson et al., 2021). However, the optimal time of the year for making the most accurate count of the taiga bean goose population remains to be investigated. Likewise, the performance of different counts has not been evaluated with data independent from the counts. Thus, it is unknown, i) whether the birds from different breeding areas are mixed with each other during the counts, ii) how spatially dispersed the population is during the counts, iii) whether birds (and how many birds) are missed in the counts, and iiii) whether birds (and how many birds) move between sites during the counts and are thereby double-counted.

We use satellite tracking data from the years 2014–2021 to study the movements and distribution of the taiga bean goose CF population during the non-breeding season. First, we describe the overall movements of the flyway population during the non-breeding season, also revealing previously unknown migration patterns. Second, we estimate the migratory connectivity of the population to reveal any clustering during the non-breeding season (and thus, whether some particular clusters could be missed in the censuses). Third, we estimate changes in the spatial distribution of the population to find the periods favourable for assessing the population size. Fourth, we compare the tracking data to the available census data from 2020–2021 to study the current performance of two different (spring and autumn) population censuses. Finally, we discuss the future perspectives to be considered when assessing population size for the taiga bean goose and other migratory populations.

## 2. Material and methods

### 2.1. Satellite tracking

We caught taiga bean geese for deployment of global positioning system (GPS) transmitter neck-collars in Denmark and Finland in the years 2014–2015 and 2018–2020, respectively. In Denmark, ten birds (all adult females) were caught using large clap nets at one wintering site, at Lille Vildmose, Jutland (56°54’N 10°13’E) by decoying wild birds with tame geese.

In Finland, 16 birds (14 females and 2 males) were caught using cannon-netting on spring staging sites at Outokumpu and Liperi (62°42’N 29°07’E) in North Karelia, and 41 birds (33 females, 8 males) were caught at the breeding sites before breeding, also using cannon-netting. These sites are located at Virrat in South Ostrobothnia (62°22’N 23°16’E), Lieksa in North Karelia (63°16’N 30°28’E), Pudasjärvi and Utajärvi in North Ostrobothnia (65°04’N 26°50’N and 65°12’N 26°52’E, respectively), and Salla in Lapland (66°51’N 28°36’E). Another two birds were caught in Lieksa (both females) and two in Utajärvi (both females) during summer when the birds were flightless due to remigial moult. For a more detailed field method description, see Piironen et al., (2021). Before the analysis, we removed two Finnish caught birds (both females) that vanished to Russia quickly after marking. Additionally, we excluded a male that was paired with another tracked bird from the analysis. Altogether, we used tracking data from 68 individuals (59 females, 9 males), which were all adults (at least two years old). For birds marked in Denmark (n = 10, all adult females), we used “Ibis” solar-powered GPS-GSM neck collars produced by Ecotone Telemetry. These transmitters weighed 30 g, which added < 1 % of the body mass of the instrumented birds. GPS resolution was set to two hours i.e. devices recorded the GPS position every second hour when battery charge levels permitted. The devices transmitted the data via the Global System for Mobile Communications (GSM) Short Message Service (SMS). Pre-deployment calibration demonstrated > 90% accuracy to within 10 m of positional data. One Danish bird caught on 14 November 2014 was followed to the Netherlands, subsequently flew to Norway but encountered severe weather and returned to Denmark, where it was retrieved dead in February 2015 (the track of which can be seen in Figure 1) and the GPS collar reused later the same year.

For birds marked in Finland (n = 58), we used OrniTrack-44 (56 birds) and OrniTrack-38 (2 birds) solar-powered GPS-GSM neckcollars produced by Ornitela UAB. OrniTrack-44 and OrniTrack-38 weigh appr. 45 and 38 grams, respectively, which added < 2 % of the weight of the body mass of the instrumented geese. These transmitters log GPS positions and send data to the server via a GSM/GPRS) network either by e-mail or SMS (short message service). To ensure the quality of the tracking data, we excluded GPS noise from the data (i.e. apparently erroneous locations such as lat 00° 00’ lon 00° 00’) and locations with hdop (horizontal dilution of precision of the GPS fix) values ≤ 2. The hdop values were only available for the OrniTrack devices.

### 2.2. Migratory connectivity and spatial distribution

We estimated the migratory connectivity of the population during the non-breeding period using Mantel’s correlation (rM), a correlation between two (distance) matrices (Cohen et al., 2017). The rM values can range between −1 and 1, so that 1 expresses full connectivity (individuals that breed close to each other are also close to each other during non-breeding season), 0 expresses no connectivity (complete mixing of population) and −1 expresses full negative connectivity (individuals breeding close to each other are far away from each other during the non-breeding season). As the origin of non-breeding geese is difficult to determine, we used only individuals with at least one breeding attempt during the tracking period (n = 42) to estimate the migratory connectivity. For those individuals, rM was calculated between the breeding site and the daily locations during the non-breeding season. For the calculation of rM, we used one location from each individual per day.

For the birds marked in Denmark (all females), we identified the nesting sites using the same method (location revisitation metrics; Picardi et al., 2020) that was previously used to identify taiga bean goose nest sites from the same tracking data (Piironen et al., 2021). However, we adjusted criteria to fit the GPS resolution (two hours) used for the birds marked in Denmark. In summary, we identified possible nest sites from the period 15 April–30 June from revisited places with the following criteria: 1) Nest site (defined as a 60-m radius to account for small-scale movements around the nest and bias in the GPS locations) must be visited on at least six consecutive days (corresponding to average clutch size and laying one egg approximately per day; Cramp & Simmons, 1977), 2) it must be visited in at least 50 % of days between first and last visit, and 3) at least 30 locations must be from the site. From the candidate nest sites, we selected the most visited site for each bird and each breeding season as the nest site (bean geese are not known to re-nest after unsuccessful attempts; Pirkola & Kalinainen, 1984). We note that these criteria include some subjective threshold values, but we believe that the conclusions about nesting based on these criteria are in accordance with what we can clearly see by following the tracks of individual birds. For birds that attempted to breed in several years, we used the centre of the different nesting sites (which were not more than a few kilometres apart from each other) as the breeding site for calculating r_M_.

Regarding birds marked in Finland, this study is based on the same satellite tracking data as the previous study by Piironen et al. (2021), so we used individual breeding sites and status provided in that study (see Additional file 2), determined using the same method as used in this study for the birds marked in Denmark. The two birds marked in 2018 in Finland were caught during moult at the breeding grounds from flocks containing adults and their offspring, and we thereby considered them as breeding birds at their breeding sites. As goose pairs move together, their movements are dependent on each other. To ensure independence of the data, we used tracking data from only one member of a goose pair to analyse the connectivity.

We estimated the spatial distribution of the population separately for each day during the non-breeding season using Minimum Convex Polygon (MCP; Mohr, 1947). We did not calculate the MCP for a period arbitrarily chosen between 1 June and 31 August, because some of the birds were marked near their breeding sites, so the choice of marking sites would affect the MCP during the breeding season. However, as the MCP is nowhere near to its minimum close to this period (Figure 2), the delineation of the excluded period is not critical for the purpose of this study i.e. for finding the optimal period for population size assessment. For the calculation of MCP, we used one location from each individual per day. To find periods when the population is the most concentrated every year (despite the variation between years), we merged the locations from each date from the years 2012–2021 before calculating the MCPs.

**FIGURE 2.**
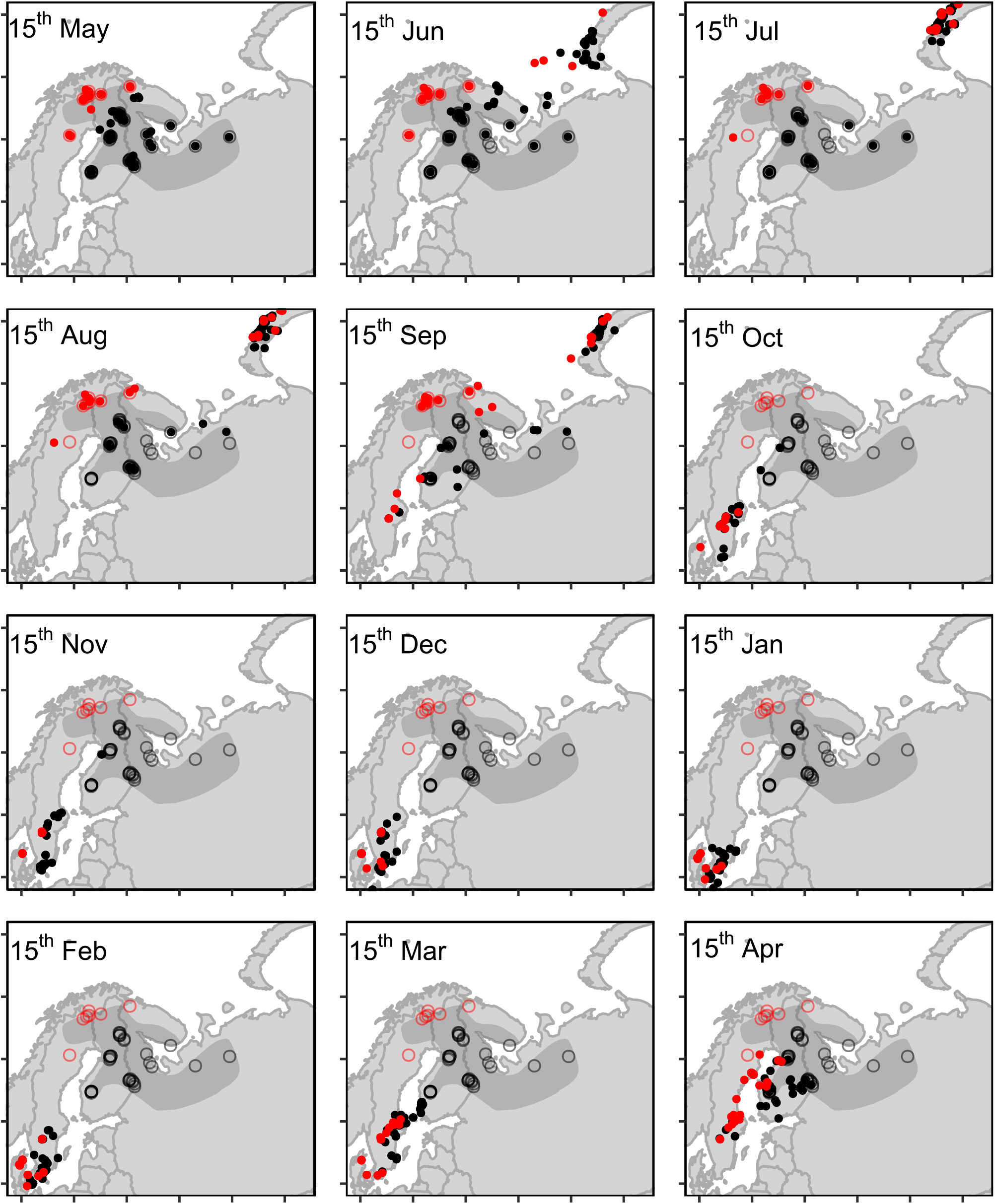
Non-breeding distribution and breeding sites of satellite-tracked taiga bean geese in 2014–2021. The non-breeding distribution is illustrated as the mid-month positions of individual birds (dots). Locations from the same date in different years are pooled to each map, i.e. each map contains one location per individual per year on a given date from the years 2014–2021. Circles denote the breeding sites for birds with at least one breeding attempt during the study period (note that the map also includes non-breeding birds, which are not connected to any of the breeding sites). Locations and breeding sites of birds marked in Finland and Denmark are illustrated with black and red, respectively. The shaded grey area denotes the breeding distribution of the Central Flyway population (redrawn after Marjakangas et al., 2015 and Heinicke et al., 2018).

We performed analysis using packages adehabitatHR (Calenge, 2006), MigConnectivity (Cohen et al., 2017) and related packages in R software version 4.1.1 (R Core Team, 2020).

### 2.3. Comparison of satellite tracking data and census data

We assessed the performance of taiga bean goose population censuses (spring and autumn) by comparing satellite tracks of tagged geese to the positions and timing of the counts from autumn 2019 (carried out on 14–25 October), spring 2020 (29 February–2 March) and spring 2021 (12–16 March). The autumn counts used in this study were carried out in addition to the standardised mid-October counts (Nilsson & Kampe-Persson, 2020). These counts are so-called total counts, i.e all birds in the population are assumed to be found and counted once, by counting birds early in the morning when they departed from the roost or later when they were feeding in the fields. The count method was selected to be suitable for different count sites. The autumn counts were carried out mainly during the morning flights, whereas the spring counts were mainly carried out at the feeding fields. The count data for autumn counts included date, time, count site (coordinates) and the number of birds counted. For the spring counts, the date is known but exact time of the day was not available. However, at the two major sites, counts were carried out during the roost flight in the morning (5.00 am – 7.00 am). At the other sites, counts were carried out during the day (9.00 am –2.00 pm) on feeding fields. For a more detailed description of the count methods, see Skyllberg & Tjernberg (2008), Kampe-Persson (2017) and Nilsson & Kampe-Persson (2020).

We compared the count data to satellite tracking data from all individuals tracked during the count (for autumn 2019 and spring 2020 n = 16, for spring 2021 n = 40). For comparison with spring count data, we used locations from the above-mentioned time intervals, as the exact time for counts was unknown. For autumn counts, we used locations from the time window of ± 30 minutes around count time (as the count time was known). Count sites in the data represent feeding areas where geese were searched for and counted (counts in the field) or the location where geese were counted during the roost flight. For the field counts, we compared the locations of satellite-tracked birds at the above-mentioned time intervals with the location of the feeding areas at which geese were counted. For roost flights, we compared the locations of the tracked birds matched with the location of the roosts, or at feeding sites close to the roost within the above-mentioned time intervals.

## 3. Results

### 3.1. Migration routes and migration phenology and migratory connectivity

The migration routes and migration phenology of satellite tracked taiga bean geese are illustrated in Figures 1 and 2. Birds marked in Denmark had breeding grounds in northern Sweden and Norway, in the Kola Peninsula and in northwestern Finland, more to the northwest than those of birds marked in Finland (Figure 2). Most of them migrated along the west coast of the Bothnian Bay unlike the birds breeding elsewhere in Finland or in Russia, which exclusively migrated through Finland east of Bothnian Bay (Figure 1).

In August, the birds were still at their breeding and moulting sites. In mid-September they began to arrive in staging areas in central Sweden, where they stayed for variable time periods until moving further south. The birds marked in Finland gathered in southern Sweden in December-February, with some individuals visiting Denmark (n = 6) and Germany (n = 2) during winter 2020–2021. The birds marked in Denmark began to arrive at the same sites for wintering in October, but note that one of these birds wintered elsewhere in Denmark (Sjælland) and one in Sweden later during the study period. The birds started to move northwards in early February, and the northward movement increased during February. In mid-March, many birds had already moved to Finland and the majority of the birds that migrate through Finland had left Sweden in mid-April. During March and April, most birds moved step-by-step to the north on either side of the Bothnian Bay, but birds heading east jumped across Finland to their breeding or staging site in eastern Finland. In mid-April, the birds were spread along their spring migration routes, as some birds were still in central Sweden while the first birds were already at their breeding sites.

### 3.2. Migratory connectivity

The strength of the migratory connectivity of the population expressed as Mantel’s correlation (rM) in the years 2019–2021 is shown in Figure 3. Among all tracked birds, connectivity stayed mainly below 0.5 in August-February, indicating moderate overall connectivity during the non-breeding season (i.e. birds from different breeding grounds do not completely mix with each other in staging and wintering areas). However, there are periods with very low connectivity (rM < 0.2), especially in the year 2021. Although there was some variation between the years, the connectivity seems to be higher during mid-winter (December-January), than during the autumn migration (September– October) or the beginning of spring migration (late February and March) in both years. Essentially, birds breeding in the northwestern breeding sites (i.e. birds marked in Denmark) show higher connectivity than the birds breeding elsewhere (i.e. birds marked in Finland). We note that this can be, to some extent, an artefact caused by the fact that all birds marked in Denmark were caught from one wintering site in north Jutland, well away from the major wintering areas in southeast Denmark. This might explain especially the high connectivity during the mid-winter, when geese were at their wintering sites (winter site fidelity is known to be high among several goose species, e.g. Wilson et al., 1991; Fox et al., 1994). However, as these birds also had somewhat separate breeding grounds (Figure 2) and more defined migration routes than birds breeding more to the east (Figure 1), there was true connectivity between the northwesternmost breeding areas and wintering areas in northern Jutland, Denmark. Nevertheless, birds from all breeding sites mixed with each other in the Swedish staging sites during the spring and autumn migration (Figure 2), which explains the lower connectivity during these periods.

**FIGURE 3.**
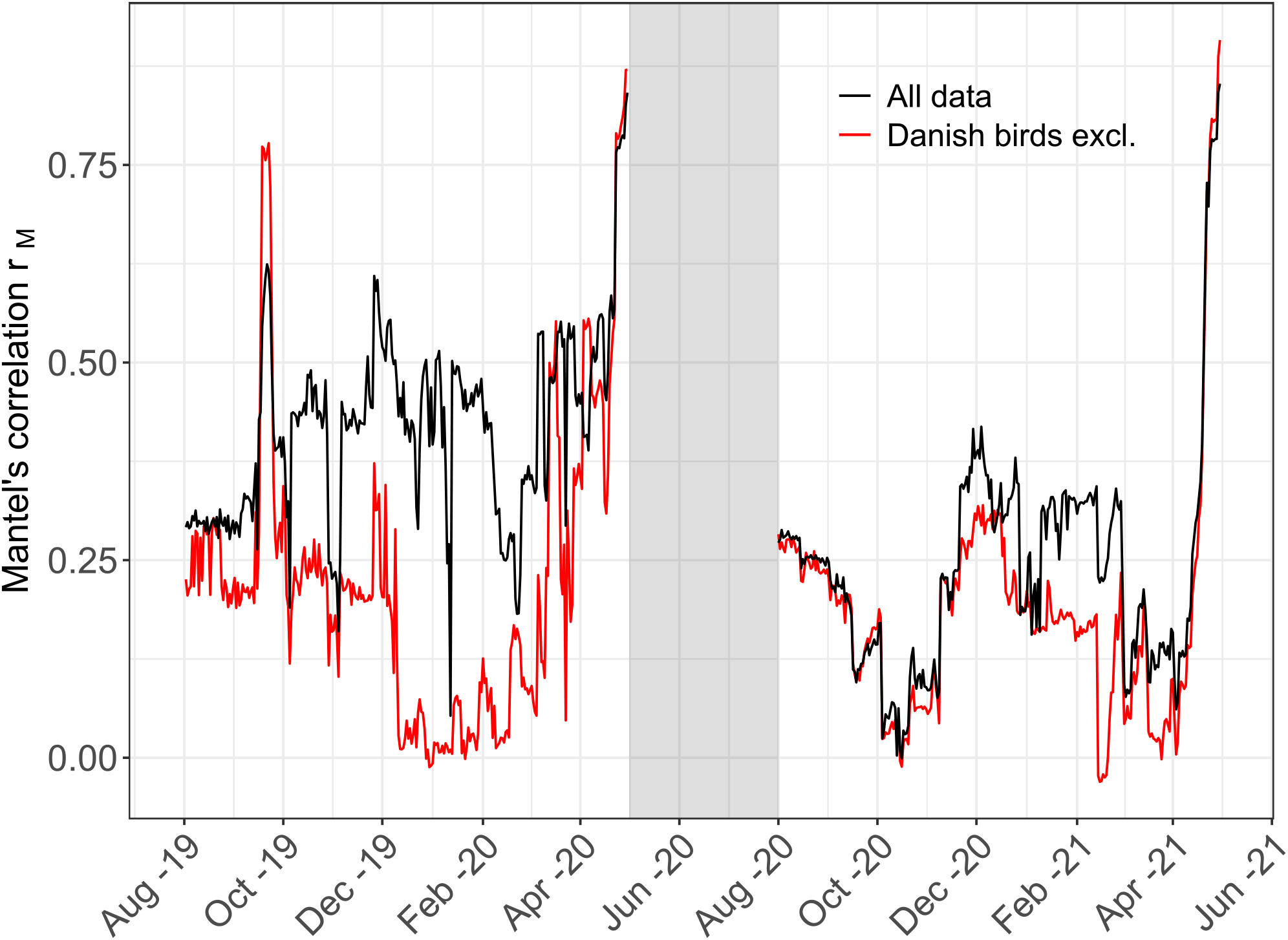
Migratory connectivity of the satellite tracked taiga bean geese during the non-breeding season from 1 August 2019 to 30 April 2021, expressed as Mantel’s correlation (rM). The shaded grey column denotes the breeding season.

### 3.3. Spatial distribution during non-breeding season

The within-year variation in the size of the area covered by the distribution of the satellite-tracked taiga bean geese is illustrated in Figure 4. In August, when the birds were still on their breeding and moulting sites, the size of the area covered by the population was relatively large. The size of the area reached its maximum in September, when the first birds moved to Sweden, while the rest of the population was still on their breeding and moulting sites (Figure 5). The remarkable reduction in the size of the area covered by the birds occurred in early October, when the birds returned from the breeding grounds in Fennoscandia and western Russia and the moulting sites in Novaya Zemlya and gathered at staging sites in central Sweden. The population was concentrated into the minimum area between late November and late December (Figure 5).

**FIGURE 4.**
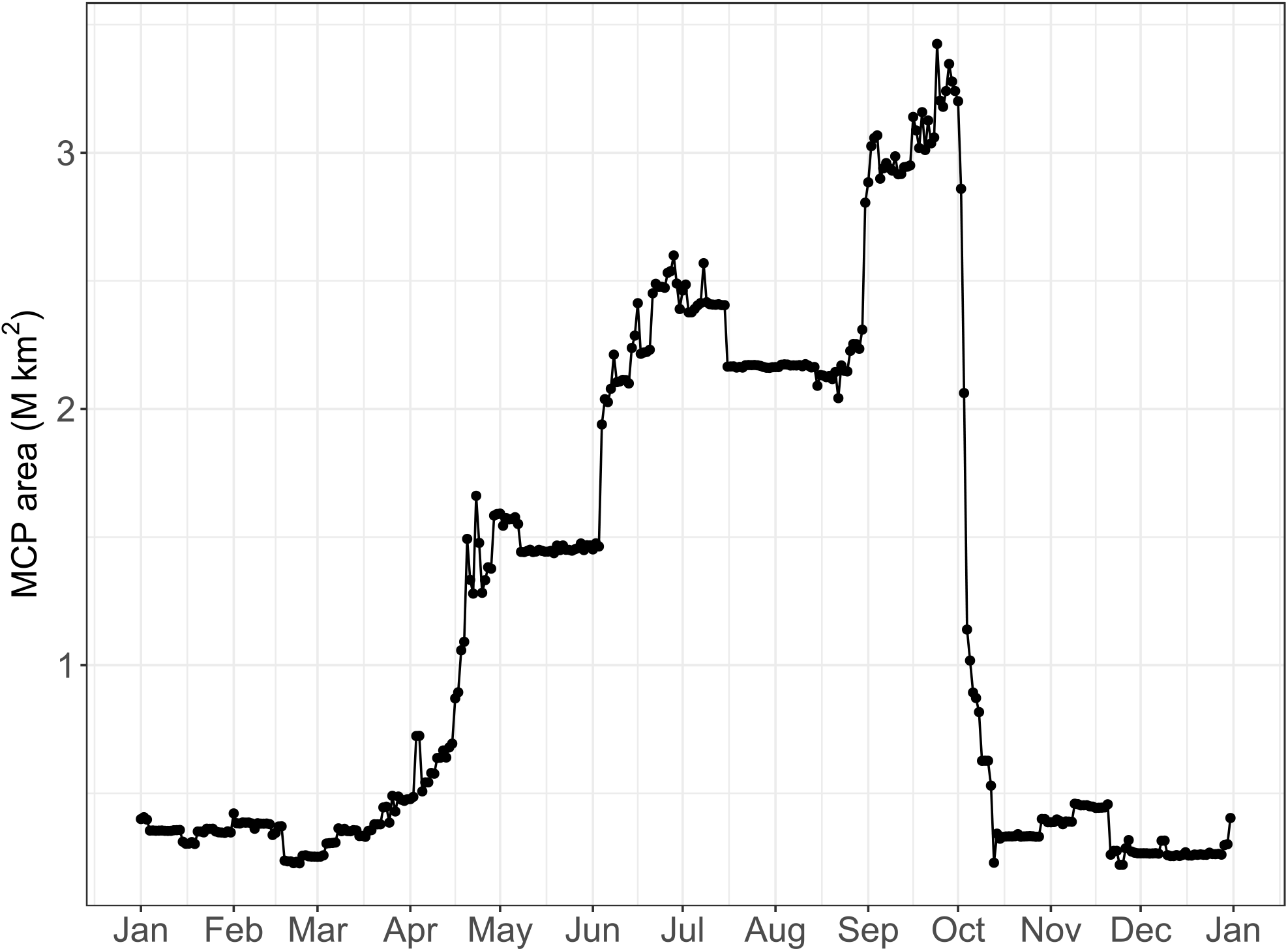
The size of the area covered by satellite tracked taiga bean geese during the non-breeding season, calculated as Minimum Convex Polygon (MCP). For the calculation of the daily MCPs, data has been merged from the years 2014–2021.

**FIGURE 5.**
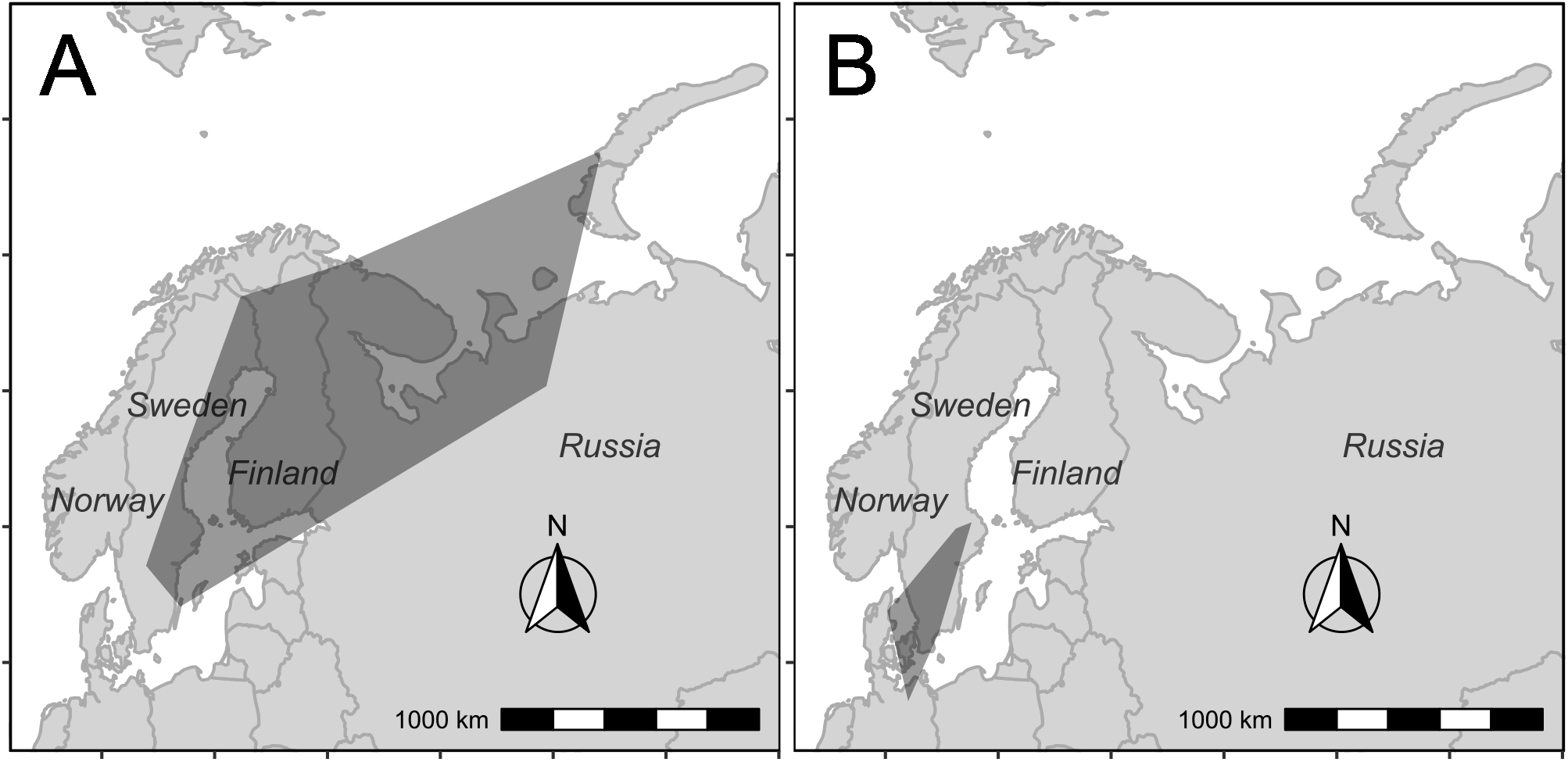
The maximum and minimum area covered by satellite tracked taiga bean geese during the non-breeding season, expressed as a Minimum Convex Polygon (MCP). Map A shows the day when MCP is at its maximum (24 September), and map B the day when MCP is at its minimum (24 November). For the calculation of the daily MCPs, data was merged from the years 2014–2021.

### 3.4. Comparison of count data and satellite tracking data

In the autumn 2019 count, six out of 16 of the satellite-tracked birds (37.5 %) were present at a count site during the count once (i.e. were on any one count site during the count). None of the birds were present on two count sites during the counts (i.e. were double-counted) and 10 birds (62.5 %) were not present at any count site during a count. In the spring count 2020, 12 out of 16 (75 %) of the birds matched with sites covered by a count once, one bird (6 %) was double-counted and three birds (19 %) were not present at any count site during a count. In the two sites where the same satellite-tracked individual was present during the counts, 3440 and 2800 birds were counted. In the count in spring 2021, 23 out of 40 (57.5 %) matched with a count once, four (10 %) were double-counted and 13 birds (32.5 %) were not near or present at count sites. Out of the total 13 birds that would not have been detected by counts, two had already migrated to Finland before the count period, four moved to Finland during the count period, and seven stayed in Sweden during the count period, but were not present at any of the count sites during the counts.

## 4. Discussion

Data from the tracked birds showed moderate to low migratory connectivity between breeding and non-breeding areas among the CF taiga bean goose population. This has consequences for population genetics as well as future research and conservation needs of the population. Both migratory connectivity and the spatial distribution (the total area instantaneously containing members of the population) of the tracked birds varied substantially within the non-breeding season, which influences the most favourable periods for population size assessment. Comparing satellite tracking and count data indicated that current autumn and spring count schemes likely underestimate true population size, even though spring and autumn counts generally exceed the corresponding winter counts (Johnson et al., 2021). Our findings provide important perspectives to be considered when studying migratory connectivity and assessing the population size of the taiga bean goose population and migratory animal populations in general.

### 4.1. Migration patterns and migratory connectivity

Our results showed that the taiga bean geese breeding in north-western Russia (Karelia, Kola Peninsula and Arkhangelsk Oblast) have similar migration patterns to the birds breeding in Finland. They migrate via Finland in autumn to winter mainly in southern Sweden, with some movements to southeast Denmark and Germany in some years (Figure 1). Our results also showed that wintering birds from northeastern Jutland in Denmark mainly breed in the westernmost parts of the taiga bean goose breeding range, the majority of which migrated along the west coast of Bothnian Bay, while some birds also migrated through Finland and breed in the Kola Peninsula and northern Finland. Despite the partially different wintering areas, all tracked birds gathered at the same staging sites in central Sweden during migrations. This decreased the strength of the migratory connectivity especially during the spring migration (Figure 3). Our results therefore confirm the recent findings of Knight et al. (2021), who showed that the connectivity can vary substantially during the annual cycle. The fact that the population can be more separated in different times of the year, can complicate population size estimation (censuses should be timed correctly to cover the whole population). It also has implications for conservation (effective actions must be focused on sites and at times when the population is most likely to be limited) and population genetics (since the population can become structured as a result of the separate timing and place of pair formation, see below). However, we require further research to reveal all implications of migratory connectivity to the conservation of migratory animal populations, not least to estimate this population metric comprehensively throughout the annual cycle.

As pair formation among waterfowl usually takes place during winter (Rohwer & Anderson, 1988), low migratory connectivity between breeding and wintering sites should lead to genetically mixed populations. Birds marked in Finland (breeding both in Finland and Russia) showed low connectivity (Figure 3), so our results are coherent with the recent study by Honka et al., (2022), who found no genetic structure among the taiga bean geese sampled in Finland. The geese wintering in Denmark showed higher migratory connectivity (Figure 3), potentially leading to genetic differentiation between the birds wintering in Denmark and Sweden. Genetic mixing among goose populations can also take place during summer on moulting grounds (as found among greater white-fronted goose *Anser albifrons*; Kölzsch et al., 2019), and taiga bean geese from the entire breeding range of the CF population have common moulting grounds in Novaya Zemlya (Piironen et al., 2021; Figure 1 in this article). Future research should concentrate on the comprehensive study of the genetic structure of taiga bean geese from different breeding origins, and on determining the timing of pair formation in taiga bean geese and its implications for the genetic structure of the population.

### 4.2. Non-breeding distribution and estimation of taiga bean goose population size

The relative size of the area including all of the tagged taiga bean geese was at its lowest from the last half of November to the beginning of January (Figure 4), implying that this is the point in the annual cycle when the population is most favourable for monitoring. The size of the area covered by the population increased slightly in the beginning of January, but remained low until mid-March, which suggests there are good reasons for continuing the current counts carried out in Sweden in mid-winter and spring. In contrast, the same results suggested that the current autumn counts (carried out in mid-October) seem vulnerable to bias caused by the fact that a part of the population remains on staging areas in Finland at that time in some years (Figure 2). The timing is also crucial with regards to the spring count, as the birds started moving northwards in February, and some birds had already arrived in Finland in early March. The correct timing will probably become even more critical in the future, especially as global warming advances the spring migration (Cotton, 2003).

Regarding the comparison between tracking data and count data, the precision of the census data (the lack of comprehensive information on the areas covered by the counts), used count methods (non-simultaneous counts) and relatively small number of satellite tracked individuals prevented us from using more sophisticated methods to assess the count data with the use of tracking data (for example, see Ganter & Madsen, 2001; Dennhardt et al., 2015; Clausen et al. 2019, Booms et al. 2021). However, the available data from these counts provided a possibility to carry out the most simple comparison between tracking and count data. Our results indicate that these counts could underestimate the true population size, as some of the tracked birds were not present in any of the count sites at the time they were counted. This is mainly caused by the birds moving between the count sites during the count period or migrating to known staging sites outside the overall count area (for example flying to Finland during the spring count). We also note that the birds from the staging sites in southwestern Finland (as well as the few birds still lingering at the wintering sites) are included in the final estimates of the taiga bean goose population size made from spring counts (Skyllberg, 2015). This is done to correct the underestimation bias caused by the birds leaving to Finland before the counts. However, it also increases the possibility for double-counting, as birds that are counted once in Sweden can be included in the bird numbers monitored at Finnish staging sites (which was the case with one satellite tracked bird in our study in the spring 2021).

To improve the current taiga bean goose censuses in the future and to increase the accuracy and transparency of the population size estimates, we suggest three actions to carry out in the future. First, the documentation of the counts should include the areas covered by the counts with precise timestamps. Second, it would be important to carry out each census simultaneously at all count sites, which would avoid some of the bias introduced by birds moving during the count (which seems to be currently the most important source of bias). Third, population size estimates (based on total counts) should be evaluated also in the future. The current tracking projects provide a starting point for the evaluations, but a greater number of tracked individuals would provide a more accurate picture on the performance of the counts.

### 4.3. Assessment of population censuses with marked individuals

Generally, our results are in line with the previous studies comparing satellite tracking data and total counts, which have revealed that total counts likely underestimate the true population sizes of various animals (Dennhardt et al., 2015; Battaile et al., 2017; Schummer et al., 2018). For geese, mark-recapture-estimates have also provided higher population size estimates than total counts (Ganter & Madsen, 2001; Clausen et al., 2019). As total counts will probably remain as a *de facto* method for assessing the size of many goose populations in the immediate future, these results confirm that performance of such counts need to be evaluated with independent data whenever possible. Although the evaluation can be done in multiple ways, the expanding usage of satellite tracking provides an easy way to study how individuals with precise locations in space and time behave in relation to the assumptions of the counts. We highlight that with a different study design than ours, it would be possible to carry out more sophisticated and rigorous evaluation of the population size assessment and form corrected population size estimates combining tracking and census data (see Dennhardt et al., 2015; Booms et al., 2021).

## Ethics statement

Capturing and marking of birds was done by the approval of Finnish Wildlife Agency (licence number 2019-5-600-01158-8) and under approval and licence to Aarhus University.

## Conflict of interest statement

None declared.

## Funding statement

This work was funded by the Ministry of Agriculture and Forestry by a grant to AP and TL (grant number 1439/03.02.02.00/2019). Finnish Wildlife Agency and Natural Resource Institute Finland (Luke) funded satellite tracking devices and field work. The October census in Sweden was financed by a grant to HKP from Stiftelsen Gustaf Adolf och Anna Maria Alvins fond till främjande av Sveriges Naturskydd and grants to Leif Nilsson from the Swedish Association for Hunting and Wildlife Management and the Swedish Environmental Protection Agency. Funding for fieldwork and purchase of tracking devices in Denmark came from the Danish Nature Agency to ORT and ADF.

## Author’s contributions

AP designed the study (together with TL), led the field work for capturing and marking of geese in Finland, ADF and ORT were responsible for the same during the earlier project in Denmark. US and HKP, provided the census data and participated in the writing of the manuscript. TL conceived the original idea, designed the study (together with AP), participated in the writing of the manuscript and supervised throughout the process. AP led the manuscript writing, aided by all other coauthors, who agreed to the final version.

## Acknowledgements

We thank numerous volunteers who participated in the bean goose counts. Additionally, we thank Matti Tolvanen, Niko Keronen, Marko Palomaa, Urpo Paavola and Juho Ristikankare for help in the fieldwork in Finland capturing birds for GPS tagging. In Denmark, we are grateful to the Å.V. Jensen Foundation and Lille Vildmose Centre for permission to catch on the site, especially to Bo Gregersen and local staff for their help and support. We express our enormous thanks to Kees Polderdijk for catching the geese, ably assisted by Jens Peder Hounisen and Michael Schmidt.

Thanks also to Thorkild Lund, Thorkil Brandt, Dorthe and Flemming Sørensen for their local knowledge and support. We acknowledge support from the Danish Ringing Centre of the National Natural History Museum and use of their rings. We also thank Leif Nilsson for collaboration regarding the mid-winter counts.

## Notes

### Competing Interest Statement

The authors have declared no competing interest.

